# Diversity of stomatal and cuticular structures affect microbial colonization in temperate forest tree species

**DOI:** 10.64898/2025.12.01.691630

**Authors:** WX Schulze, ED Schulze, S Reiße, R Rischke, O Bouriaud, B Büdel, T Straub, E Pillai, B Tanunchai, W Purahong, S Simm, M Noll

**Affiliations:** Department of Plant Systems Biology, University of Hohenheim, 70599 Stuttgart, Germany; Max-Planck-Institute for Biogeochemistry, Jena, Germany; Core Facility Hohenheim, University of Hohenheim, 70599 Stuttgart, Germany; Department of Applied Natural Sciences and Health, Coburg University of Applied Sciences and Arts, Coburg, Germany; Stefan cel Mare University of Suceava, Suceava, Romania; RPTU Kaiserslautern, Department of Biology, Kaiserslautern, Germany; Department of Soil Science, Helmholtz Centre for Environmental Research, Halle, Germany

**Keywords:** Surface structural features, microscopic imaging, broad leaf, coniferous needle, stomata, microbiome

## Abstract

Throughout their life cycle, tree leaves are subject to colonization and degradation by microorganisms, including fungi, bacteria, and algae. These relationships co-evolved with chemical properties, leaf shape, and surface structures. Here we quantified leaf abaxial surface texture features based on variables extracted from scanning electron microscopic images, resulting in an abstract quantitative texture complexity score. This complexity score was used to test functional hypotheses such as growth habitat preferences and microbial colonization patterns. Here we show that the evolution of leaf surface texture complexity traits correlated with anatomical features such as stomatal density and leaf orientation and correlated with Ellenberg temperature habitat indicator. Furthermore, increasing leaf surface texture complexity was found to be negatively correlated with plant pathogen richness (broad-leaved species) or lichenization (conifers), suggesting protection effects. Moreover, we found a negative correlation of leaf surface texture complexity with fungal and bacterial specialists. Our results highlight leaf surface texture complexity as a key, previously underappreciated trait shaping microbial diversity and leaf-microbe interaction patterns. This opens promising avenues for future research on plant-microbe co-evolution, trait-based ecosystem modeling, and the potential use of surface traits in forest management and disease resistance strategies.

## Introduction

The textural complexity of leaves in vascular plants has fascinated botanists since the early use of Scanning Electron Microscopy (SEM) [1, 2]. Plant scientists have used these advances in microscopy to study evolution, function and patterning of the stomata, which regulate gas exchange between the leaf interior and the atmosphere [3]. At the same time, leaves and needles offer a variety of habitats for microorganisms, including bacteria, fungi [4], and algae [5], which colonize the leaves from bud break until leaf senescence [6]. Boosted by advances in sequencing technologies, investigations of the “jungle” of microbial and algal communities that inhabit leaf surfaces [5, 7] have become an important research focus of plant and microbial science in recent years.

Stomatal and cuticular features are likely co-evolutionary adaptations shaped by respective growth habitats. This process is believed to have begun for gymnosperms approx. 298 million years ago, and for angiosperms approx. 200 million years ago [3]. The colonization of leaf surfaces by microbial communities is also presumed to have evolved in a host- and environment-specific manner [8]. Consequently, leaf surface structures may have become key determinants in shaping these organismic interactions. Generally, the ecological roles and functional traits of microorganisms on and within fresh leaves range from pathogenic to mutualistic [9], and fungi are able to shift their ecological roles depending on habitat conditions [6]. Fungal epi- and endophytes were suggested to be the primary saprotrophs following leaf senescence and abscission [4, 9].

Interestingly, a variety of bacteria, oomycetes, and fungi exploit stomatal openings as major invasion routes [10-12] primarily due to the availability of moisture and possibly inorganic ions. To prevent microbe invasion, plants recognize the so-called microbe-associated molecular patterns (MAMPs) which are highly conserved within a class of microbes, such as flagellin for bacteria and chitin oligosaccharides for fungi. The detection of these molecules then triggers stomatal immunity and defense responses, including stomatal closure or inhibition of stomatal opening [13-15]. In turn, some plant pathogens have devised various strategies to manipulate stomata behaviour, facilitating easier invasion [12, 16].

While the composition of microbial communities on leaf surfaces was studied intensively [17], the role of structural, chemical, and elemental leaf surface features has received less attention, although they provide important factors driving colonization [18]. Moreover, methods of implementing leaf surface structures on microscopic level as quantitative variables were developed only recently and concepts to quantify and classify species by the features of surface structures were not yet widely exploited under consideration of functional feedbacks.

On a macroscopic scale of forest landscapes, structural features, such as tree diameter, tree height, dead wood volume, diversity of bark types, were shown to have high explanatory power for the species richness, especially for plants and some insect groups [19]. On the scale of individual organs, the biochemical properties of leaves, such as reflectance and transmittance spectra were recently used to classify leaf phenotypes for cultivar classification and selection during breeding [20]. Interestingly, the leaf spectral emissions are also affected also by leaf surface properties [21], However, the lack of sufficiently large-scale quantitative data on leaf epidermal surface characteristics has so far slowed the progress in this research direction [22]. Another classification approach utilized epidermal texture patterning by extracting texture features from microscopic images. These features enabled quantitative structural comparisons, which have led to a significant increase in correct classification of plant species [23]. The classifications based on preprocessed images of leaf surface offer the opportunity for higher throughput compared to traditional morphometry studies using transversal leaf cuts [24, 25]. Recently, microscopic leaf surface analysis has been further developed for taxonomic identification of plants [26], focusing primarily on four image features such as shape, texture, vein patterns, and color. The advantage of such leaf texture classification lies in its ability to exclude noise during segmentation and feature extraction, which can result in over 90% correct species classification [23]. Thus, species classification based on texture traits of leaf surfaces bears great potential for correlative testing of functional hypotheses.

Here, we used machine learning methods to extract features from microscopic images of abaxial leaf epidermal surface and developed a ranking of texture complexity based on game-theoretic approaches [27, 28]. The development of a quantitative complexity measure added explainability to species classification and enabled testing of functional hypotheses regarding the role of leaf epidermal surface texture complexity in dominant tree species of a European temperate forest. In the following, we hypothesize that leaf surface texture complexity is correlated to evolutionary drivers, such as leaf orientation and habitat preference. Here we propose that leaf epidermal surface texture complexity may be an important determinant of the diversity of microbial communities and other organismic interactions, which in turn feed back into leaf functioning.

## Materials and Methods

### Study site and sampling

The mature leaves and needles were collected from a privately managed mixed temperate forest in the Hainich-Dün region of Thuringia, Germany (51°120N 10°180E). The forest site is growing at about 500 m elevation with 600 to 800 mm precipitation and 6 to 7.5 °C mean annual temperature. The bedrock is limestone with 30 to 1 m of cambisol soil mainly from weathering (see also Tanunchai et al., 2022). Briefly, leaves and needles (Supplementary table S1) were sampled from the ground up to 6 m at the sun-exposed southern edge of a tree crown using a telescopic leaf cutter. Most of the sampled tree species were collected in a single stand of about 5 ha. Several leaves of up to five trees per species were collected; sampling was conducted in batches throughout the season. For conifers, fresh needles from the current growing season were sampled. Thus, we examined a forest community where trees interact at the canopy level to resolve leaf surface structures independent of variation in site conditions. For microbiome analysis, a minimum of 200 g leaves and needles per tree individual were collected (each with five true replicates (tree)) in October 2019, with new gloves in sterilized plastic bags. The samples were transported on ice within 3 h to the laboratory and frozen at -80°C before further processing. For microscopy, leaves were collected from the same site in October 2024. Leaves were analyzed freshly, but also after freezing them as part of storage at −80°C. All leaf samples were analyzed together.

### Scanning Environmental Microscopy (SEM)

Surface structures of the leaves were examined using a scanning electron microscope (EVO15, Carl Zeiss Microscopy GmbH). Fresh leaf material from 5 to 20 different leaves was analyzed without fixation or coating with conductive materials, and under conditions of significantly reduced chamber vacuum (610 Pa). Images were collected at two different magnification scales (1900x / 230x). Additionally, the instrument is equipped with an Energy Dispersive X-ray Spectroscopy (EDS) detector (UltimMax, Oxford Instruments), enabling elemental analysis alongside and overlaying the imaging. The distribution of C, N, and O in the surface structure was determined for selected species. Element distribution was then quantified using AZtec software Version 6.0 (Oxford Instruments).

### Confocal microscopy

Cell walls of hyphae were stained by Calcofluor White and analyzed by confocal microscopy (Zeiss LSM 900). Calcofluor White stains beta-1,3 and beta-1,4 polysaccharides, which interact with chitin or peptidoglycan structures [29, 30]. Each needle or leaf section was incubated in the staining solution (Calcofluor White diluted 1:1 with 10% KOH, [vol/vol]) for 20 min in the dark, followed by thorough washing with water. A laser with a wavelength of 405 nm was used to excite the Calcofluor White-colored structures and the fluorescence was detected in the emission range from 405 to 460 nm. The autofluorescence of the chlorophyll was generated with a laser with a wavelength of 640 nm and detected in the range from 640 nm to 700 nm. Images were processed using the LSM Plus deconvolution algorithm and analyzed and exported with the ZEN 3.6 blue edition software (Carl Zeiss Microscopy GmbH).

### Quantification and classification of leaf surface texture complexity

SEM images were used as databases to quantify the texture complexity of leaf abaxial surfaces. For analysis of leaf structural complexity, images from leaf areas without fungal colonization were used. All images from all species were analyzed together. The primary stages of our analysis include sub-image sampling, feature computation, classification of species, and complexity ranking. For sub-image sampling, a fine-tuned model, Detectron2 [31], was applied for precise segmentation of stomata within the SEM images. This targeted approach helped in isolating key features, such as structural variations of wax formation, relevant to the texture complexity analysis. We then randomly sampled 120 sub-images around the identified stomata within a larger image, each of size ca. 80 x 80 µm^2^. Sample images were resized to 512 x 512 pixel dimensions to ensure uniformity in feature extraction and analysis. Balancing the number of samples per species results in a dataset of 12960 images. Texture features were computed for each image using the Gray-Level Co-occurrence Matrix (GLCM), spectral features using Fourier transform techniques, combined with Principal Component Analysis (PCA) to distill the images down to 25 significant spectral features, gradient-based attributes, and the compressed size of the image files (Supplementary Figure S1A). The latter provides insights into the inherent texture complexity of the image data due to the lossless coding. For classification of leaf surfaces from tree species based on the computed features, we utilized the k-nearest neighbors (kNN) algorithm with k=3. This method has been chosen based on preliminary tests indicating its effectiveness for our dataset and the performance matched previously reported results [23].

Ranking of leaf surface texture complexity features from different species was achieved by a game-based approach (Supplementary Figure S1B), which was used for its versatility, scalability and the possibility of simple interpretation and an easily performed all-against-all comparison. This game utilized a set of interpretable and uncorrelated features (image compression size, ASM, and gradient standard deviation), each contributing uniquely to the perceived complexity of a leaf. In the game, individual leaf samples were pitted against each other in a pairwise comparison format. During each match-up, the sample exhibiting greater texture complexity, as determined by our set of defined features, was awarded a point, while the other sample lost a point. This dynamic scoring method ensured that the contribution of each feature to overall texture complexity was considered, allowing for a nuanced ranking of leaf/needle surface texture complexity on species-level. We applied the Bradley-Terry model [27] to compute a single score for each species. Species were grouped into distinct classes using hierarchical clustering based on a pairwise comparison with the Kruskal-Wallis test.

### Leaf parameters

Stomatal density was calculated from the SEM images. Leaf angle was measured in the field using an inclinometer. Ellenberg habitat indicator values were obtained from a database [32]. Nutrient contents of leaves from the sampled trees were already published previously [33]. Other tree parameters, such as rooting depth, rooting type and mycorrhization listed in Supplementary Table S2 were derived from literature [34, 35].

### DNA extraction and Illumina sequencing

Detailed methods on DNA extraction and Illumina MiSeq Sequencing of mature leaves and needles were previously described elsewhere [6, 33]. Briefly, we initially subsampled up to 10 leaves or needles per tree, which were then washed with sterile Tween solution (0.1% vol/vol) and thereafter deionized water. Thereafter samples were ground by using liquid nitrogen and a pestle, homogenized, and stored at −20 °C. For DNA extraction, approximately 120 mg of homogenized leaves and needles were processed using the DNeasy PowerSoil Kit (Qiagen, Hilden, Germany) following the manufacturer’s instructions. Fungal internal transcribed spacer region (ITS)-based amplicon sequencing using the Illumina MiSeq platform was carried out as described previously [36].

### Bioinformatics

We re-analyzed an existing ITS sequence dataset obtained on a subset of the twelve tree species (8 broad-leaf, 4 conifers; [6]). The same trees were sampled for sequencing (in previous work) and for complexity analysis (this study). The ITS rDNA sequences were processed to amplicon sequence variants (ASVs) as explained earlier [6]. Ultimately, 2451 rarefied fungal ASVs were obtained, each with a minimum sequencing depth of 21,967 sequences per sample. The fungal ecological function of each ASV was assessed using FungalTraits [37] following the authors’ guidelines. The classification of generalists and specialists of a taxonomic table was calculated based on the deviation of niche width indices (Shannon, Levins, or occurrence) from null values computed with permutation algorithms for community matrices using the spec.gen function in the EcolUtils package in R (https://github.com/GuillemSalazer/EcolUtils [38]). We classified the leaf colonizing fungi and bacteria as specialists when they were found established on only one target tree species, as generalists when found common on all tree species, or as opportunists growing selectively on conifers or broad-leaved trees.

## Results

We followed a hypothesis-driven correlative approach, systematically quantifying the contribution of whole leaf traits to leaf surface texture complexity of 21 broad-leaved tree species and six needle-leaved conifers (Table S1; Figure 1A). The texture complexity of leaf surface was tested for its dependence on leaf morphological features and habitat preferences (Supplementary Table S2) and for its contribution to interactions with other organisms and their feedback to tree function. We hypothesize that genetic determinants of leaf morphology influence the leaf surface structural complexity and that this complexity, in turn, affects interactions with other organisms. In this study, we tested the effects of leaf surface structural complexity on interactions with microorganisms.

**Figure 1:**
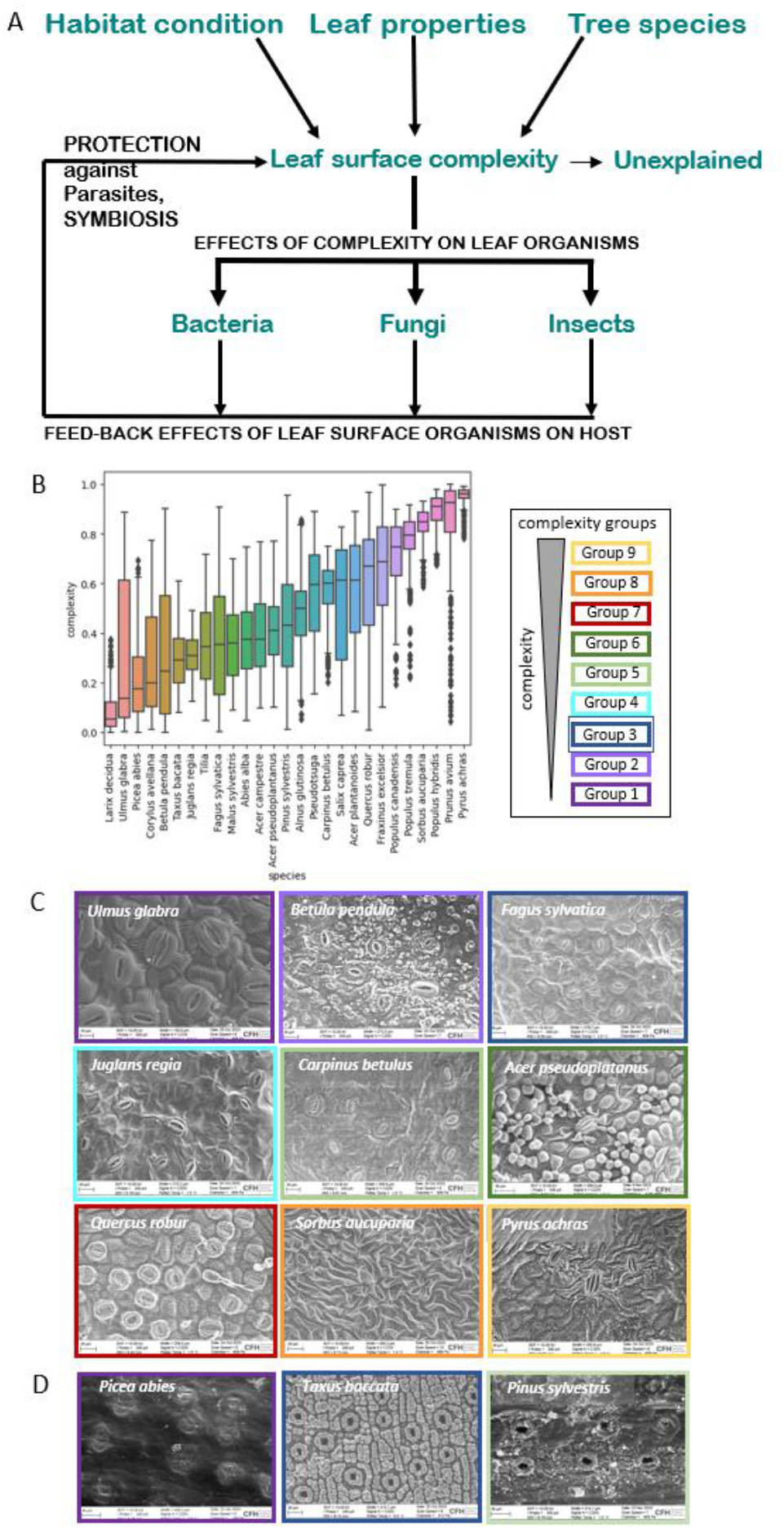
Texture complexity of abaxial leaf side across 27 tree species. (**A**) Experimental design based on hypotheses tested regarding tree and habitat characteristics affecting surface texture complexity. Leaf texture complexity in turn may affect colonization by various organisms, which may then feed back on tree function. (**B**) Ranking of texture complexity scores across sampled species. Complexity groups were defined by ANOVA hierarchical clustering. (**C**) Images of representative broad-leaved species within the nine complexity groups. (**D**) Images of representative coniferous species from different complexity groups.

### Development of a quantitative surface structure complexity score

The surface texture complexity was derived as a comparative value from grayscale texture patterns on scanning electron microscopy images of the leaf abaxial side (Supplementary Figure S1) and resulted in quantitative relative ranking of the 27 tree species from low texture complexity (*Ulmus glabra, Fagus sylvatica* and *Larix decidua*) to high texture complexity (*Sorbus aucuparia* and *Prunus avium*) (Figure 1B, C, D). The resulting ranking of leaf surface texture complexity showed high accuracy (96%) in separating tree species. Variance in surface texture complexity (low variance: *Sorbus aucuparia* and *Prunus avium, Pyrus achras*; high variance *Ulmus glabra, Betula pendula*, and *Faxinus excelsior*) was mainly due to stochastic occurrence of leaf surface features, such as the spatial organization of stomata, the stomatal di-morphism (e. g. *Prunus avium*), and other structural components like trichomes or glands. To our surprise, 60% of the broad-leaved species have dimorphic stomata (i.e. stomata of different size), most pronounced in *Prunus avium*. Dimorphic stomata were located on veins and originated from earlier development in the process of leaf expansion, where veins are formed prior to the mesophyll [39]. The calculated texture complexity values were robust against independent ranking runs, and also when limiting the calculation to the stomata as area of interest near stomata (Supplementary Figure S2). To assess batch-to-batch variations (e.g. from leaves collected throughout the season), for three species (*Acer pseuoplatanus, Fagus sylvatica, Quercus robur*) we calculated separate complexity ranks for each of the respective batches (Supplementary Figure S2). The complexity ranking of individual batches was within total variation for the respective species. Thus, in the following, we used the complexity ranking for each species involving all images from the whole leaf area averaged across all leaf samples.

### Abaxial leaf surface texture complexity may contribute to leaf surface microenvironment

Tree taxonomy did not strictly relate to the observed leaf surface texture complexity (Supplementary Figure S3), i.e. closely related species can display highly different surface structure patterns (e.g. *Acer* species). In broad-leaved species, leaf surface complexity showed a positive correlation with leaf angle (r^2^=0.42, p=0.02): leaves with a more horizontal angle showed lower leaf surface complexity. By contrast, in conifers, stomatal density (r^2^=0.85, p=0.02), but not stomata pore size showed a positive correlation with leaf surface complexity (Supplementary Figure S4A). Interestingly, Ellenberg temperature habitat indicator [32], but not light indicator or moisture indicator showed a considerable correlation with leaf texture complexity in broad-leaved species as well as in conifers (r^2^=0.36, p=0.08 for broad-leaved species; r^2^=0.95, p=0.16 for conifers; Supplementary Figure S4B). This suggests that also epidermal surface features in addition to tree architecture and leaf anatomy, may affect the microenvironment of the leaf and thus the preferred growth habitat. Leaf temperature in turn has been shown to be a major factor for pathogens [40].

### Abaxial leaf surface texture complexity protects against pathogens and lichen formation

Many fungi, algae, and cyanobacteria can colonize leaves and needles as pathogens, saprophytes, symbionts or in the form of lichens. We used an existing data set of leaf microbial communities [6] on a subset of twelve tree species (8 broad-leaf, 4 conifers) to test for interdependence of microbial communities with leaf surface properties (Table S3). Samples for the analysis of microbial communities were taken from the same trees as samples for leaf surface complexity analysis. Total richness of fungi was not affected by leaf surface texture complexity (Supplementary Figure S5A). However, we observed a negative correlation (r^2^=0.88, p=0.02) of the total number of colonizing bacteria with leaf surface texture complexity of broad-leaved species (Supplementary Figure S5B). Across all tree species, and for fungi as well as bacteria, higher leaf texture complexity significantly (p<0.01) resulted in lower richness in specialists (Figure 2A), but not in generalists (Supplementary Figure S5C). This relationship was largely driven by the significantly higher richness of specialists on coniferous species, especially *Picea*.

**Figure 2:**
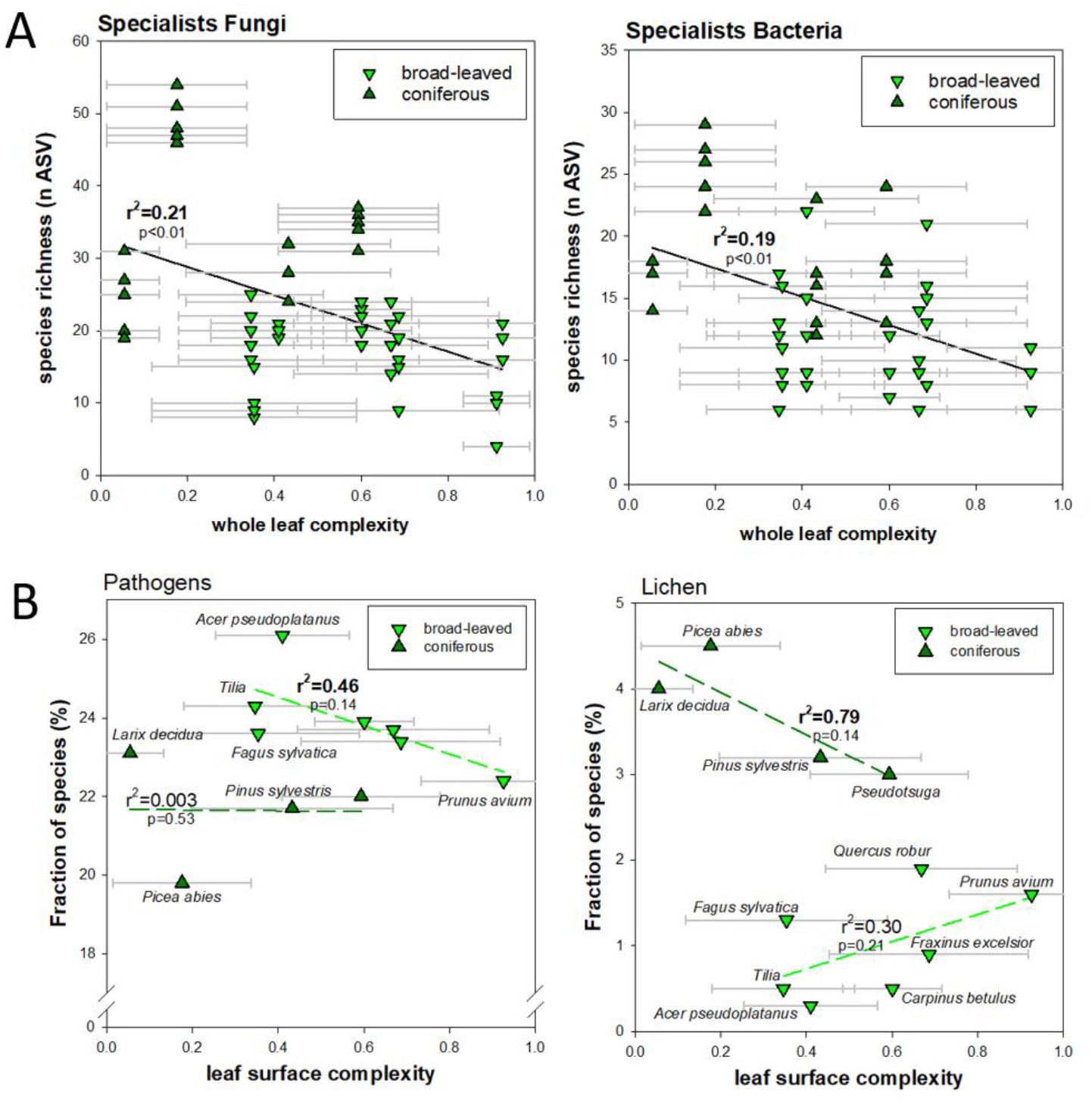
Relationship of fungal and bacterial richness with leaf surface texture complexity. (**A**) Correlation of richness in fungal and bacterial specialists with leaf surface complexity. (**B**) Correlation of the proportion of fungal pathogens or lichen-based fungi with leaf surface texture complexity. Data points represent averages with standard deviation. Species richness was assessed by amplified sequence variants (ASVs).

When separating fungal species by their ecotypes, we observed a considerable negative correlation of the proportion of pathogens with leaf surface texture complexity in broad-leaved species (Figure 2B). Thus, our data suggests leaves with higher surface texture complexity had a tendency for less pathogens (r^2^=0.46, p=0.14). For conifers, we observed a negative correlation of the proportion of lichen colonization (r^2^=0.79, p=0.14) with leaf surface texture complexity (Figure 2B). We propose that complex leaf epidermal surface structures present an important means of protection against pathogens (broad-leaved) or heavy lichenization (conifers). The interaction of leaf surfaces with leaf colonizing fungi and lichens was therefore investigated in more detail also with respect to nutrient sources for the microbiome.

### Leaf surface is nutrient poor and requires organismic interactions for epiphyllic growth

Abaxial leaves and needles were found rich in covalently bound carbon (C) and oxygen (O) molecules but were poor in nitrogen (N) (Supplementary Figure S6). The element distribution was linked mainly to structural components: the stomatal pores themselves were more O-rich compared to guard cell surfaces (e. g. *Quercus*, Supplementary Figure S6A). The atomic composition in needles from *Abies alba* shifted over time, reflecting a temporal succession (2, 4, and 8 years of needles on the tree) of co-occurring fungal hyphae, which were visible by their more O-rich surface chemistry compared to the needle surface wax layer (Supplementary Figure S6B to D). Indeed, fungal hyphae showed higher N content compared to the needle background, reflecting chitinous cell walls of fungi and/or glycan molecules in N-fixing cyanobacteria. Epiphyllic fungi were frequently found associated with algae and cyanobacteria (Figure 3). Such observed partnerships can reflect different organismic interaction types, ranging from parasitic interactions by phagocytosis (Figure 3A), mutualistic by aggregation (Figure 3B,C,D), or symbiotic by encapsulation of cyanobacteria or algae (Figure 3E,F,G), representing initial stages of lichenization. Initiating at the epistomatal wax layers of the stomatal antechamber, the fungal structures over time formed dense networks (Figure 3H), which in the form of lichens ultimately covered whole branches (Figure 3I).

**Figure 3:**
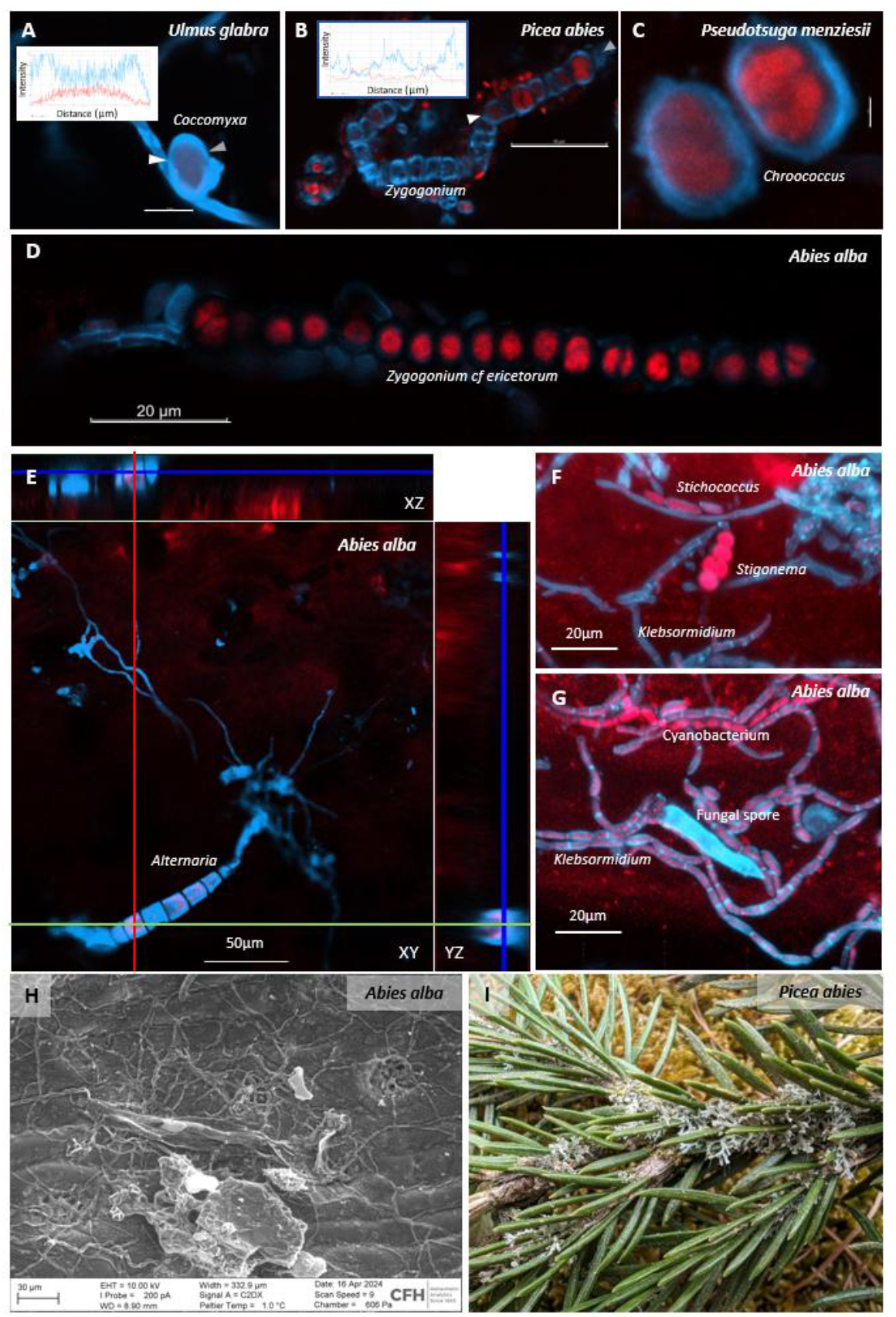
Association of fungi with algae and cyanobacteria. (**A**) Phagocytosis of the algae *Coccomyxa*. Scale bar 5µm. (**B**) Threads of the green alga *Zygogonium*. Scale bar 50µm. (**C**) Unicellular cyanobacteria. *Chroococcus* Scale bar 2µm. (**D**) Threads of green algae Zygogonium. Scale bar 20µm. (**E**) Fungal hyphae and spores of *Alternaria* form a layer above the epidermal cells as indicated by XYZ ortho projections. Scale bar 50µm. (**F**) Cyanobacterium *Stigonema* as well as green algae *Stichococcus* and *Klebsormidium*. Scale bar 20 µm.(**G**) Network of algae, cyanobacteria and fungi. Scale bar 20 µm. (**H**) Hyphae form a network between epistomatal wax plates. Scale bar 100 µm. (**I**) *Anaptychia ciliaris* (dominant), *Candelariella xanthostigma* (yellow spots, rare species), *Melanohalea exasperatula* (brown-green layer), and *Apatococcus lobatus* (green algae) growing on 3-year-old needles of *Picea abies* (identification by Burkhard Büdel). In A to G, red shows chlorophyll autofluorescence, cyan shows chitin or peptidoglycan in cell walls stained by Calcofluor White (LSM900, Carl Zeiss Microscopy GmbH). Inserts show fluorescence intensity profiles of calcofluor white (cyan) and chlorophyll (red) along a line between white and gray arrows.

The source of nutrient acquisition of foliicolous lichens from host leaves remains unclear [41-43]. Apart from the N produced by the cyanobiont through N_2_-fixation, mineral N from host leaves or needles has been suggested as an alternative or additional source of nutrients for the lichens. Mineral N (N_Mn_) and dissolved organic C (DOC) are water-soluble nutrient sources immediately available for surface-associated microbes. These nutrients can originate from atmospheric deposition (e.g. nitrate, ammonium; [44]), by a molecular film of liquid water that also allows ion transport across the cuticle [45], or by exploiting plant apoplasmic space after deterioration of the cuticle (Figure 4). The C-assimilation is driven by photosynthesis in the photobiont, which is highly dependent on water, light, and bioavailable N [46]. The differences in water potential between lichen thallus and host substrate, as well as within the lichen symbiont, provide a suitable environment for nutrient transport [47]. We used regression models to analyze the contribution of the leaf surface texture complexity and leaf nutrient content in influencing the fungal and bacterial leaf surface communities (Table 1). Leachable total organic C, organic N, the N-to-P ratio, and Ca concentration in host leaves were significantly correlated with both fungal and bacterial community composition, indicating their influence on the colonizing microbiome. In coniferous trees, P concentration was significantly correlated with the fungal community, while Mg concentration was significantly correlated with the bacterial community (Table 1). Interestingly, in conifers, leaf surface complexity values calculated around regions of interest (ROI) at stomata showed an R^2^ value 0.74 (p=0.052) in the correlation with bacterial communities, suggesting the contribution of surface texture complexity to bacterial community composition was in a similar magnitude as TOC or magnesium. For fungal community composition, highest contribution of leaf surface complexity was found across all tree species (R2=0.42/0.46; p-value=0.091/0.066). We conclude that while nutrient composition of the leaf turned out as important factor defining the leaf microbiome, the surface texture complexity turned out as additional component not to be neglected, especially in conifers.

**Table 1.** Coefficients of determination statistics of environmental variables fitted to non-metric multidimensional scaling (NMDS) ordination of lichenized fungal community composition based on relative abundance data and nutrient contents from previous work [33]. The analysis was performed using the vegan package in R. Bold letters indicate statistically significant influences of the respective feature.

| Fungi | *All Tree (n = 12)* |  | *Broad leaf (n = 8)* |  | *Needle (n = 4)* |  |
| --- | --- | --- | --- | --- | --- | --- |
| | $R^2$ | $P$ | $R^2$ | $P$ | $R^2$ | $P$ |
| Complexity | 0.4265 | 0.091 | 0.0603 | 0.9583 | 0.2735 | 0.515 |
| ROI complexity | 0.4640 | 0.066 | 0.0499 | 0.9583 | 0.2862 | 0.473 |
| TOC | **0.6974** | **0.009** | **0.9847** | 0.2083 | 0.1435 | 0.705 |
| Nmin | 0.2852 | 0.193 | **0.6247** | 0.6250 | 0.1130 | 0.677 |
| Norg | **0.6167** | **0.009** | **0.6768** | 0.5417 | 0.2028 | 0.496 |
| C to N ratio | 0.0681 | 0.743 | **0.9864** | 0.2917 | **0.5331** | 0.108 |
| C to P ratio | 0.3161 | 0.187 | **0.6076** | 0.6667 | **0.5781** | 0.127 |
| N to P ratio | **0.6471** | **0.009** | **0.7947** | 0.6667 | **0.6538** | **0.048** |
| Ca | **0.6724** | **0.006** | **0.8078** | 0.5000 | 0.4999 | 0.125 |
| Fe | **0.5178** | **0.035** | **0.9960** | 0.0833 | 0.2030 | 0.523 |
| K | 0.3925 | 0.117 | 0.2479 | 0.8750 | **0.7009** | 0.111 |
| Mg | 0.3261 | 0.185 | **0.8325** | 0.3750 | 0.2187 | 0.444 |
| P | 0.4665 | 0.060 | 0.4783 | 0.7500 | **0.7477** | **0.031** |

| Bacteria | *All Tree (n = 12)* |  | *Broad leaf (n = 8)* |  | *Needle (n = 4)* |  |
| --- | --- | --- | --- | --- | --- | --- |
| | $R^2$ | $P$ | $R^2$ | $P$ | $R^2$ | $P$ |
| Complexity | 0.2904 | 0.251 | 0.3570 | 0.8750 | **0.5904** | 0.156 |
| ROI complexity | 0.3406 | 0.196 | 0.4023 | 0.8333 | **0.7472** | 0.052 |
| TOC | **0.8545** | **0.001** | **0.9979** | 0.0417 | **0.7503** | **0.045** |
| Nmin | 0.4189 | 0.114 | **0.8288** | 0.4583 | 0.2901 | 0.515 |
| Norg | **0.7803** | **0.002** | **0.7493** | 0.5833 | 0.3289 | 0.451 |
| C to N ratio | 0.2719 | 0.286 | **0.9837** | 0.0417 | 0.2510 | 0.571 |
| C to P ratio | 0.2937 | 0.230 | 0.4950 | 0.7500 | 0.2004 | 0.672 |
| N to P ratio | **0.7821** | **0.005** | **0.8045** | 0.3333 | 0.3050 | 0.517 |
| Ca | **0.6872** | **0.007** | **0.9815** | 0.1667 | 0.0541 | 0.903 |
| Fe | 0.4900 | 0.075 | **0.8214** | 0.5000 | 0.1994 | 0.659 |
| K | 0.0816 | 0.699 | 0.2724 | 0.8333 | 0.2810 | 0.478 |
| Mg | **0.5984** | 0.036 | **0.6925** | 0.6667 | **0.7811** | **0.034** |
| P | 0.4609 | 0.094 | 0.3908 | 0.7917 | 0.0983 | 0.821 |

**Figure 4:**
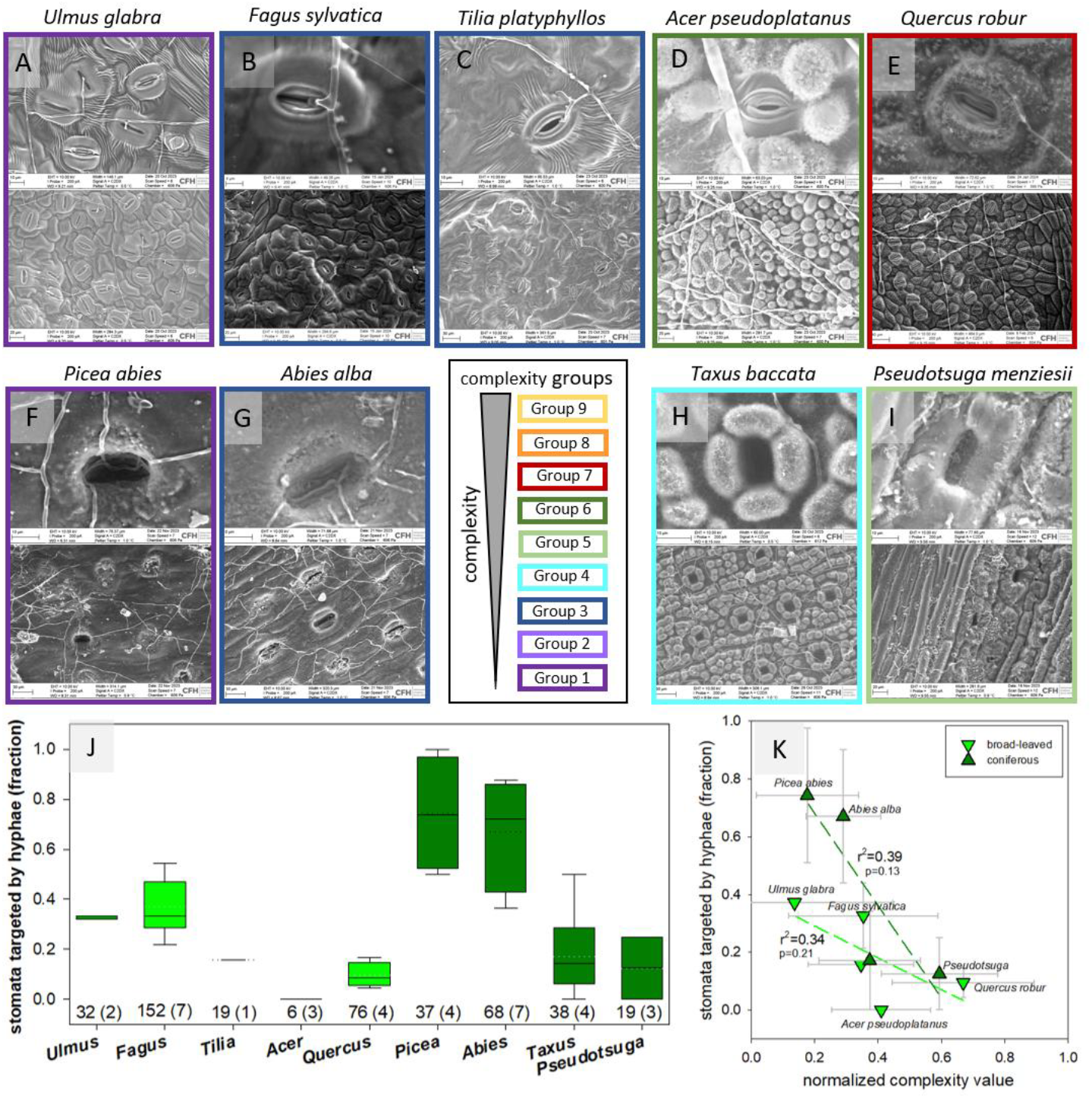
Fungal hyphae growth is directed towards stomata. Broad-leaved species: (**A**) *Fagus sylvatica*, (**B**) *Ulmus glabra*, (**C**) *Tilia platyphyllos* from different texture complexity groups. Protective structures around stomata prevent fungal penetration in (**D**) *Acer pseudoplatanus*, (**E**) *Quercus robur*. Coniferous species: (**F**) *Picea abies*, (**G**) *Abies alba* stomata are heavily targeted by hyphae, while in (**H**) *Taxus baccata* and (**I**) *Pseudotsuga menizesii* protective structures were observed. Images were acquired on a scanning electron microscope (EVO15, Carl Zeiss Microscopy GmbH), border colors indicate respective ANOVA complexity groups. Representative examples are given at closeup magnification of single stomata and an overview of lower magnification. Border colors indicate respective ANOVA complexity groups. (**J**) Fraction of stomata targeted by fungal hyphae. Box plot shows the scatter of data (solid line: median; dotted line: mean; whiskers: 5th/95^th^ percentile). The number of evaluated stomata is shown, with the number of evaluated images in brackets. For conifers, images of one year old needles were used. (**K**) Relationship between the fraction of stomata targeted by hyphae and leaf surface texture complexity. Data are shown as averages with standard deviation.

### Stomata as targets of fungal growth and initiation of lichen formation

Most fungal colonizers organized their proliferation towards stomata, irrespective of whether tree species were broad-leaved or coniferous (Figure 4A to I). Moreover, considerably higher frequency of stomatal invasions was observed on leaves with lower leaf surface texture complexity score (Figure 4J, K). Interestingly, in species with dimorphic stomata, the large stomatal pores were not observed to be penetrated by fungal hyphae. We therefore conclude that leaf surface texture complexity plays an important role in protecting stomata structures from access by fungal colonizers. This relationship was found more pronounced in the needles of coniferous trees (Figure 4K), in which microbial colonization clearly initiated at the stomatal antechambers.

Apparently, the leaf surface contains structures, for which functions in regulation of leaf microenvironment (Supplementary Figure S4) or in protection against (pathogenic) colonization (Figure 4) could be shown.

## Discussion

Our work provides evidence that microscopic structures on the lower leaf surface significantly affect the organismic interactions with leaf-colonizing microbes, especially in the context of defense against plant pathogens. These relationships became evident through the machine learning supported texture complexity analysis of microscope images developed in this study, which allowed quantitation and ranking of leaf epidermal surface complexity traits.

### Determination and ranking of leaf surface texture complexity

Texture, as defined as a statistical distribution of gray tones, has also become increasingly used in plant sciences for the classification of image features. Since texture depends on spatial scale, smooth textures at larger scales become rough as the scale decreases, e. g. in microscope images [25]. This can be overcome by computing gray-level co-occurrence matrices (GLCM) from which typical texture features such as energy, entropy, contrast, absolute value, inverse difference, and homogeneity can be extracted and used as feature vectors for machine learning models. Based on isotropic GLCM, a 90% accuracy can be achieved for plant species classification [23, 26]. For denoising and decreasing the inter-class variation, typically wavelet and fractal dimensions are used during the extraction of texture features [48]. This successfully increased the prediction accuracy of plant species based on leaf surface patterns [23].

Based on these existing methods of feature extraction from images, our workflow was built on four steps: sub-image sampling, feature computation, species classification, and complexity ranking (details see Methods), and overall it resulted in a classification accuracy of 96%, aligning with previously reported results [23]. We chose our training-validation process to be aware of intra-class variation [49] by balancing our dataset through use of multiple images of different areas per individual leaf in addition to different leaves. The addition of the game theory approach allowed the quantitative ranking of different tree species by their leaf surface texture complexity values in an all against all comparison. Major advantages of applying the game theoretical approach were its versatility, scalability and the possibility of simple interpretation. Additionally, by treating each extracted segment as an individual during the ranking of the complexity traits, the game theory approach added to high intra-class robustness. The resulting quantitative ranking of leaf surface texture complexity values was used as a basis for analysis of functional hypotheses regarding whole plant abiotic and biotic parameters. In some species (*Betula, Fagus*) we observed a larger within-species variation of leaf surface texture complexity compared to other species (*Juglans, Prunus*). This may partly be attributed to different numbers of images available for each species, but further research is needed to carefully assess within-tree variation, for example in sun or shade leaves.

### Leaf surface texture complexity and environmental factors

A previous hypothesis for micro-scale cuticular surface structure proposed that uneven structures avoid the formation of water films [1]. Liquid water covering the stomata could significantly reduce the gas exchange of leaves through the stomata. In consequence, the exchange of CO_2_ with the atmosphere would be impaired and affect growth and survival of trees. The avoidance of water films and the associated establishment of epiphyllous microbial films of organisms is crucial particularly for the upper adaxial surface of leaves in terms of light harvest. However, here we analyzed the lower, abaxial surface of leaves, which in contrast to the upper leaf surface contains the stomata and which get fully wet only rarely, depending on leaf orientation. Interestingly, we observed a positive correlation of leaf surface complexity with leaf angle but also with temperature habitat indicator (Supplementary Figure S4). We therefore propose that leaf surface texture complexity indeed contributes to leaf microenvironments regarding leaf moisture and temperature. It emerges that leaf surface structure may not as much regulate transpiration directly, but rather affect microbial community establishment and the efficiency of pathogen entry, possibly through a component of affecting leaf surface temperature properties [40] and surface water availability.

### Stomata as entry pores for fungi

We frequently observed stomata to be penetrated by hyphae and stomatal antechambers as initiation points for microbial colonization. At this stage, it remains unknown how fungal hyphae detect the location of stomata. The total amount of water vapor emitted by stomatal cells exceeds the flux of CO_2_ by about a factor 100 [50], suggesting moisture as a major determinant for stomata directed growth of hyphae. In addition, a large range of volatile organic substances are emitted at lower concentrations.

It has previously been suggested that fungi use resources excreted by the epidermal cells [7, 17]. However, the outer cell wall of epidermal cells is sealed by waxes of variable chemistry [18], and epidermal cells do not contain chloroplasts. Alternatively, it was proposed that cell walls of the stomatal cells are covered by a molecular film of liquid water that also allows ion transport across the cuticle [45]. We scanned the elemental composition of the leaf surfaces regarding C, O and N (Supplementary Figure S6) and could not detect any evidence for major nutrient leaching suggesting that epiphyllic organisms are required to be either symbiotic with cyanobacteria and algae, saprophytic or pathogenic.

Interestingly, growth and differentiation of *Uromyces appendiculatus*, a stomata-penetrating fungus infecting conifers (*Pinus*), was affected by surface topography, specifically by physical ridges of at least 0.5 µm. These minute barriers prevented fungal hyphae of finding the stomatal pore [51] supporting our findings of lower pathogenic growth on leaves of higher structural complexity (Figure 4). We can expect also other fungi to be able to distinguish unique patterns in the leaf surface during the colonization process, suggesting that indeed surface features could have defensive functions. Moreover, increased fungal competition for nutrients as well as host defense mechanisms could lead to higher numbers of specialists, which was indeed observed for species with lower leaf surface texture complexity. Given that microbial community composition was available for only 12 out of the 27 the species for which leaf surface complexity was analyzed, these barely significant relationships between leaf surface texture complexity and leaf microbe community (Figures 2, Table 1) are remarkable and require further attention in future studies.

It will remain an effort of future research to precisely identify the fungal species that are affected by structural barriers, and to separate fungal and bacterial functional groups in a modeling approach.

Furthermore, despite there being a huge analytical effort, in future it will be desirable to increase statistical power by increasing the number of observations, i.e. the number of species for which epiphyllic microbiomes were studied.

## Conclusions

We took a machine-learning based approach to quantify and rank epidermal surface texture complexity traits of European tree species. This approach of texture complexity quantitation can be generalized to other biological systems and processes. We found that surface texture complexity is an important trait in defending against microbial colonization, especially pathogens. The occupation of the lower leaf surface may be detrimental to the leaf even though the organisms are not parasitic by growing into and blocking stomatal chambers. Although our study is observational in nature, it offers a new framework for understanding the role of surface morphology in plant-microbe interactions. Future experimental work will be essential to disentangle causal mechanisms. Nevertheless, this approach opens new avenues for exploring the ecological and evolutionary relevance of microscopic surface traits, with potential implications for biodiversity, ecosystem resilience, and tree health management in a changing climate.

## Data availability

Environmental Scanning-Electron Microscopy images used for texture complexity ranking and code for complexity quantification were deposited at GitHub (https://github.com/Waltraud-Schulze/leaf-surface-complexity).

Training data are available at figshare (https://doi.org/10.6084/m9.figshare.28377767.v1).

## Acknowledgements

We thank Daniel Thomas, Bayreuth Center of Ecology and Environmental Research (BayCEER), Bayreuth, Germany for fruitful discussions on the topic. We acknowledge the contacts in the cost action “CA22158 – Exploiting Plant-Microbiomes Networks and Synthetic Communities to improve Crops Fitness (MiCropBiomes)”.

## Author contributions

WXS: analyzed data, wrote the manuscript

EDS: generated data, analyzed data, wrote the manuscript

SR: generated data, analyzed data, wrote sections of the manuscript

RR: analyzed data, wrote sections of the manuscript

OB: analyzed data, wrote sections of the manuscript BB: analyzed data

TS: generated data, analyzed data

EP: analyzed data

BT: generated data, analyzed data WP: generated data

SS: analyzed data, wrote sections of the manuscript

MN: generated data, analyzed data, wrote the manuscript

## References

1. Bresinsky A, Körner C, Kadereit JW, Neuhaus G, Sonnewald U. Lehrbuch der Botanik. 36 ed. Heidelberg: Spektrum Verlag; 2008.

2. Troughton J, Donaldson LA. Probing Plant Structure: A Scanning Electron Microscope Study of Some Anatomical Features in Plants and the Relationship of These Structures to Physiological Processes. University of Michigan: McGraw-Hill; 1972. 116 p.

3. Clark JW, Harris BJ, Hetherington AJ, Hurtado-Castano N, Brench RA, Casson S, et al. The origin and evolution of stomata. Current biology : CB. 2022;32(11):R539–R53. doi: 10.1016/j.cub.2022.04.040. PubMed PMID: 35671732.

4. Tanunchai B, Ji L, Schroeter SA, Wahdan SFM, Larpkern P, Lehnert AS, et al. A poisoned apple: First insights into community assembly and networks of the fungal pathobiome of healthy-looking senescing leaves of temperate trees in mixed forest ecosystem. Frontiers in plant science. 2022;13:968218. doi: 10.3389/fpls.2022.968218. PubMed PMID: 36407586; PubMed Central PMCID: PMC9669904.

5. Büdel B, Friel T, Beyschlag W. Biology of algae, lichens and bryophytes. Stuttgart: Springer; 2024.

6. Tanunchai B, Ji L, Schroeter SA, Wahdan SFM, Hossen S, Delelegn Y, et al. FungalTraits vs. FUNGuild: Comparison of Ecological Functional Assignments of Leaf- and Needle-Associated Fungi Across 12 Temperate Tree Species. Microbial ecology. 2023;85(2):411–28. doi: 10.1007/s00248-022-01973-2. PubMed PMID: 35124727; PubMed Central PMCID: PMC9958157.

7. Vacher C, Hampe A, Porte AJ, Sauer U, Compant S, Morris CE. The Phyllosphere: Microbial Jungle at the Plant-Climate Interface. Annual Review of Ecology, Evolution and Systematics. 2016;47:1–24.

8. Mesny F, Hacquard S, Thomma BP. Co-evolution within the plant holobiont drives host performance. EMBO reports. 2023;24(9):e57455. doi: 10.15252/embr.202357455. PubMed PMID: 37471099; PubMed Central PMCID: PMC10481671.

9. Koivusaari P, Tejesvi MV, Tolkkinen M, Markkola A, Mykra H, Pirttila AM. Fungi Originating From Tree Leaves Contribute to Fungal Diversity of Litter in Streams. Frontiers in microbiology. 2019;10:651. doi: 10.3389/fmicb.2019.00651. PubMed PMID: 31001228; PubMed Central PMCID: PMC6454979.

10. Grimmer MK, John Foulkes M, Paveley ND. Foliar pathogenesis and plant water relations: a review. Journal of experimental botany. 2012;63(12):4321–31. doi: 10.1093/jxb/ers143. PubMed PMID: 22664583.

11. Lin PA, Chen Y, Ponce G, Acevedo FE, Lynch JP, Anderson CT, et al. Stomata-mediated interactions between plants, herbivores, and the environment. Trends in plant science. 2022;27(3):287–300. doi: 10.1016/j.tplants.2021.08.017. PubMed PMID: 34580024.

12. Melotto M, Underwood W, He SY. Role of stomata in plant innate immunity and foliar bacterial diseases. Annual review of phytopathology. 2008;46:101–22. doi: 10.1146/annurev.phyto.121107.104959. PubMed PMID: 18422426; PubMed Central PMCID: PMC2613263.

13. Arnaud D, Hwang I. A sophisticated network of signaling pathways regulates stomatal defenses to bacterial pathogens. Molecular plant. 2015;8(4):566–81. doi: 10.1016/j.molp.2014.10.012. PubMed PMID: 25661059.

14. Melotto M, Zhang L, Oblessuc PR, He SY. Stomatal Defense a Decade Later. Plant physiology. 2017;174(2):561–71. doi: 10.1104/pp.16.01853. PubMed PMID: 28341769; PubMed Central PMCID: PMC5462020.

15. Ye W, Murata Y. Microbe Associated Molecular Pattern Signaling in Guard Cells. Frontiers in plant science. 2016;7:583. doi: 10.3389/fpls.2016.00583. PubMed PMID: 27200056; PubMed Central PMCID: PMC4855242.

16. Gudesblat GE, Torres PS, Vojnov AA. Stomata and pathogens: Warfare at the gates. Plant signaling & behavior. 2009;4(12):1114–6. doi: 10.4161/psb.4.12.10062. PubMed PMID: 20514224; PubMed Central PMCID: PMC2819434.

17. Vorholt J. Microbial life in the phyllosphere. Nature Reviews in Microbiology. 2012;10:828–40.

18. Riederer M, Müller C. Biology of plant cuticles. Oxford: Blackwell; 2006.

19. Storch F, Boch S, Gossner MM, Feldhaar H, Ammer C, Shall P, et al. Linking structure and species richness to suport forest biodiversity monitoring at large scale. Annals of Forest Science. 2023;80(3):1–17. doi: 10.1186/s13595-022-01169-1.

20. Peters RD, Noble SD. Characterization of leaf surface phenotypes based on light interaction. Plant methods. 2023;19(1):26. doi: 10.1186/s13007-023-01004-2. PubMed PMID: 36932424; PubMed Central PMCID: PMC10024457.

21. Holmes MG, Keiller DR. Effects of pubescence and waxes on the reflectance of leaves in the ultraviolet and photosynthetic wavebands: a comparison of a range of species. Plant, cell & environment. 2002;25(1):85– 93.

22. Baranoski GV. Modeling the interaction of infrared radiation (750-2500 nm) bith bifacial and unifacial plant leaves. Remote Sens Environ. 2006;3:335–47.

23. da Silva NR, Oliveira M, Filho HAA, Pinheiro LFS, Rossatto DR, Kolb RM, et al. Leaf epidermis images for robust identification of plants. Scientific reports. 2016;6:25994. doi: 10.1038/srep25994. PubMed PMID: 27217018; PubMed Central PMCID: PMC4877573.

24. Junior JJdMS, Backer AR, Rossatto DR, Kolb RM, Bruno OM. Measuring and analyzing color and texture information in anatomical leaf cross sections: an approach using computer vision to aid plant species identification. Botany. 2011;89(7).

25. Ramos E, Fernandez D. Classification of leaf epidermis microphotographs using texture features. Ecological Informatics. 2009;4(3):177–81.

26. Sachar S, Kumar A. Survey of feature extraction and classification techniques to identify plant through leaves. Expert Systems With Applications. 2021;167:114181. doi: 10.1016/j.eswa.2020.114181.

27. Bradley RA, Terry ME. Rank analysis of incomplete block designs. I. The method of paired comparions. Biometrika. 1952;39:324–45.

28. Huang TK, Weng RC, Lin C-J. Generalized Bradley-Terry Models and Multi-Class Probability Estimates. Jurnal of Machne Learning Research. 2006;7:85–115.

29. Henriques BS, Garcia ES, Azambuja P, Genta FA. Determination of Chitin Content in Insects: An Alternate Method Based on Calcofluor Staining. Frontiers in physiology. 2020;11:117. doi: 10.3389/fphys.2020.00117. PubMed PMID: 32132935; PubMed Central PMCID: PMC7040371.

30. Rasconi S, Jobard M, Jouve L, Sime-Ngando T. Use of calcofluor white for detection, identification, and quantification of phytoplanktonic fungal parasites. Appl Environ Microbiol. 2009;75(8):2545–53. doi: 10.1128/AEM.02211-08. PubMed PMID: 19233954; PubMed Central PMCID: PMC2675195.

31. Wu Y, Kirillow A, Massa F, Lo W-Y, Girshick R. Detectron2 2019. Available from: https://github.com/facebookresearch/detectron2.

32. Tichy L, Axmanova I, Dengler J, Guarino R, Jansen F, Midolo G, et al. Ellenberg-type indicator values for European vascular plant species. Journal of Vegetation Science. 2022;34:e13168. doi: 10.1111/jvs.13168.

33. Tanunchai B, Schroeter SA, Ji L, Wahdan SFM, Hossen S, Lehnert AS, et al. More than you can see: Unraveling the ecology and biodiversity of lichenized fungi associated with leaves and needles of 12 temperate tree species using high-throughput sequencing. Frontiers in microbiology. 2022;13:907531. doi: 10.3389/fmicb.2022.907531. PubMed PMID: 36187953; PubMed Central PMCID: PMC9523249.

34. Wang B, Qiu YL. Phylogenetic distribution and evolution of mycorrhizas in land plants. Mycorrhiza. 2006;16(5):299–363. doi: 10.1007/s00572-005-0033-6. PubMed PMID: 16845554.

35. Köstler JN, Brückner E, Bibelriether H. Die Wurzeln der Wald-bäume: Paul Parey; 1968.

36. Weißenbecker C, Schnabel B, Heintz-Buschart A. Dadasnake, a Snakemake implementation of DADA2 to process amplicon sequencing data for microbial ecology. GigaScience. 2020;9(12):giaa135.

37. Pölme S, Abarenkov K, Nilsson RH, Lindahl BD, Clemmensen KE, Kauserud H, et al. FungalTraits: a user-friendly traits database of fungi and fungus-like stramenopiles. Fungal Diversity. 2021;105:1–16.

38. Salazar G. EcolUtils: Utilities for community ecology analysis 2023. Available from: https://github.com/GuillemSalazer/EcolUtils.

39. Napp-Zinn K. Anatomie des Blattes. II. Anatomie der Angiospermen. Berlin: Gebrüder Bornträger; 1973.

40. Bernard F, Sache I, Suffert F, Chelle M. The development of a foliar fungal pathogen does react to leaf temperature! The New phytologist. 2013;198(1):232–40. doi: 10.1111/nph.12134. PubMed PMID: 23373986.

41. Grimm M, Grube M, Schiefelbein U, Zuhlke D, Bernhardt J, Riedel K. The Lichens’ Microbiota, Still a Mystery? Frontiers in microbiology. 2021;12:623839. doi: 10.3389/fmicb.2021.623839. PubMed PMID: 33859626; PubMed Central PMCID: PMC8042158.

42. Grube M, Cardinale M, de Castro JV, Jr., Muller H, Berg G. Species-specific structural and functional diversity of bacterial communities in lichen symbioses. ISME J. 2009;3(9):1105–15. doi: 10.1038/ismej.2009.63. PubMed PMID: 19554038.

43. Liba CM, Ferrara FI, Manfio GP, Fantinatti-Garboggini F, Albuquerque RC, Pavan C, et al. Nitrogen-fixing chemo-organotrophic bacteria isolated from cyanobacteria-deprived lichens and their ability to solubilize phosphate and to release amino acids and phytohormones. Journal of applied microbiology. 2006;101(5):1076–86. doi: 10.1111/j.1365-2672.2006.03010.x. PubMed PMID: 17040231.

44. Schulze ED. Air Pollution and Forest Decline in a Spruce (Picea abies) Forest. Science. 1989;244(4906):776–83. doi: 10.1126/science.244.4906.776. PubMed PMID: 17802236.

45. Burkhardt J, Basi S, Pariyar S, Hunsche M. Stomatal penetration by aqueous solutions--an update involving leaf surface particles. The New phytologist. 2012;196(3):774–87. doi: 10.1111/j.1469-8137.2012.04307.x. PubMed PMID: 22985197.

46. Palmqvist K. Tansley Review No. 117: Carbon economy in lichens. The New phytologist. 2000;148(1):11–36. doi: 10.1046/j.1469-8137.2000.00732.x. PubMed PMID: 33863029.

47. Potkay A, Ten Veldhuis MC, Fan Y, Mattos CRC, Ananyev G, Dismukes GC. Water and vapor transport in algal-fungal lichen: Modeling constrained by laboratory experiments, an application for Flavoparmelia caperata. Plant, cell & environment. 2020;43(4):945–64. doi: 10.1111/pce.13690. PubMed PMID: 31759337.

48. Tao Y, Lam ECM, Tang YY. Feature extraction using wavelet and fractal. Pattern Recognition Letters. 2001;22(3-4):271–87. doi: 10.1016/S0167-8655(01)00003-4.

49. Naresh YG, Nagendraswamy S. A novel fuzzy LBP based symbolic representation technique for classification of medicinal plants. 3rd IAPR Asian Conference on Pattern Recognition (ACPR); Kuala Lumbur, Malaysia 2015.

50. Schulze ED, Beck E, Buchmann N, Clemens S, Müller-Hohenstein K, Scherer-Lorenzen M. Plant Ecology. Stuttgart: Springer; 2009. 926 p.

51. Hoch CH, Staples RC, Whitehead B, Comeau J, Wolf ED. Signaling for Growth Orientation and Cell Differentiation by Surface Topography in Uromyces. Science. 1987;235:1659–62.

