## Supplementary Materials for "Diversity of stomatal and cuticular structures affect microbial colonization in temperate forest tree species"

### Supplementary Tables:

**Table S1**: List of species analyzed by SEM.

**Table S2**: Parameters of leaf surface features and texture complexity values

**Table S3**: Species identified based on amplified sequence variants from (Tanunchai *et al.*, 2023).

**Table S4**: Ecosystem fluxes at different study sites of broad-leaved, coniferous and mixed forests with different dominant tree species.

### References in Supplementary Materials

**Durka W, Michalski SG**. 2012. Daphne: a dated phylogeny of a large European flora for phylogenetically informed ecological analyses. *Ecology* **93**, 2297.

**Tanunchai B, Ji L, Schroeter SA, Wahdan SFM, Hossen S, Delelegn Y, Buscot F, Lehnert AS, Alves EG, Hilke I, Gleixner G, Schulze ED, Noll M, Purahong W**. 2023. FungalTraits vs. FUNGuild: Comparison of Ecological Functional Assignments of Leaf- and Needle-Associated Fungi Across 12 Temperate Tree Species. *Microb Ecol* **85**, 411-428.

**Tichy L, Axmanova I, Dengler J, Guarino R, Jansen F, Midolo G, Nobis MP, Van Meerbeek K, Acic S, Attorre F, Bergmeier E, Biurrun I, Bonari G, Bruelheide H, Campos JA, Carni A, Chiarucci A, Cuk M, Custerevska R, Didukh Y, Dite D, Dite Z, Dziuba T, Fanelli G, Fernandez-Pascual E, Garbolino E, Gavilan RG, Gegout J-C, Graf U, Güler B, Hajek M, Hennekens SM, Jandt U, Jaskova A, B. J-A, Julve P, Kambach S, Karger DN, Karrer G, Kavgaci A, Knollova I, Kuzemko A, Küzmic F, Landucci F, Lengyel A, J. L, Marceno C, Moeslund JE, Novak P, Perez-Haase A, Peterka T, Pielech R, Pignatti A, Rasomavicius V, Rusina S, Saatkamp A, Silc U, Skvorc Z, Theurillat J-P, Wohlgemuth T, Chytry M**. 2022. Ellenberg-type indicator values for European vascular plant species. *Journal of Vegetation Science* **34**, e13168.

**Supplementary Figures:**

**Supplementary Figure S1: Determination of Leaf Surface Texture Complexity.** (**A**) Workflow of quantification of leaf surface texture complexity as described in Materials and Methods. (**B**) Game theoretic approach for constructing the complexity scores.

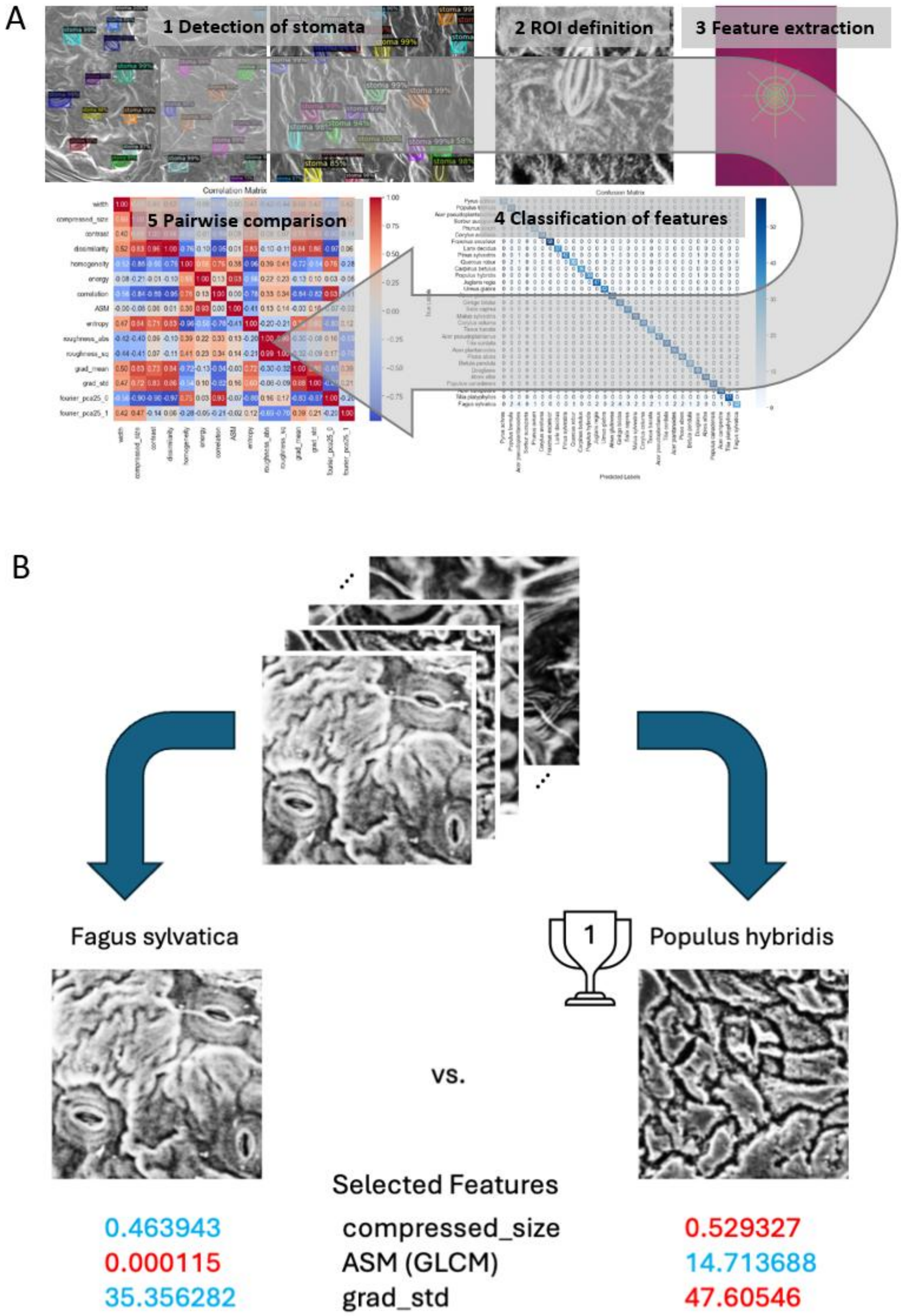

**Supplementary Figure S2: Variation of complexity ranking.** Complexity value of the different species as calculated independently based on image samples from the whole leaf (complexity 1, complexity 2) or focused on the stomata area (complexity ROI). Complexity ranking was performed separately for two different batches of leaves (yellow sqares), which was available for *Acer pseudoplatanus*, *Fagus sylvatica*, and *Quercus robur*).

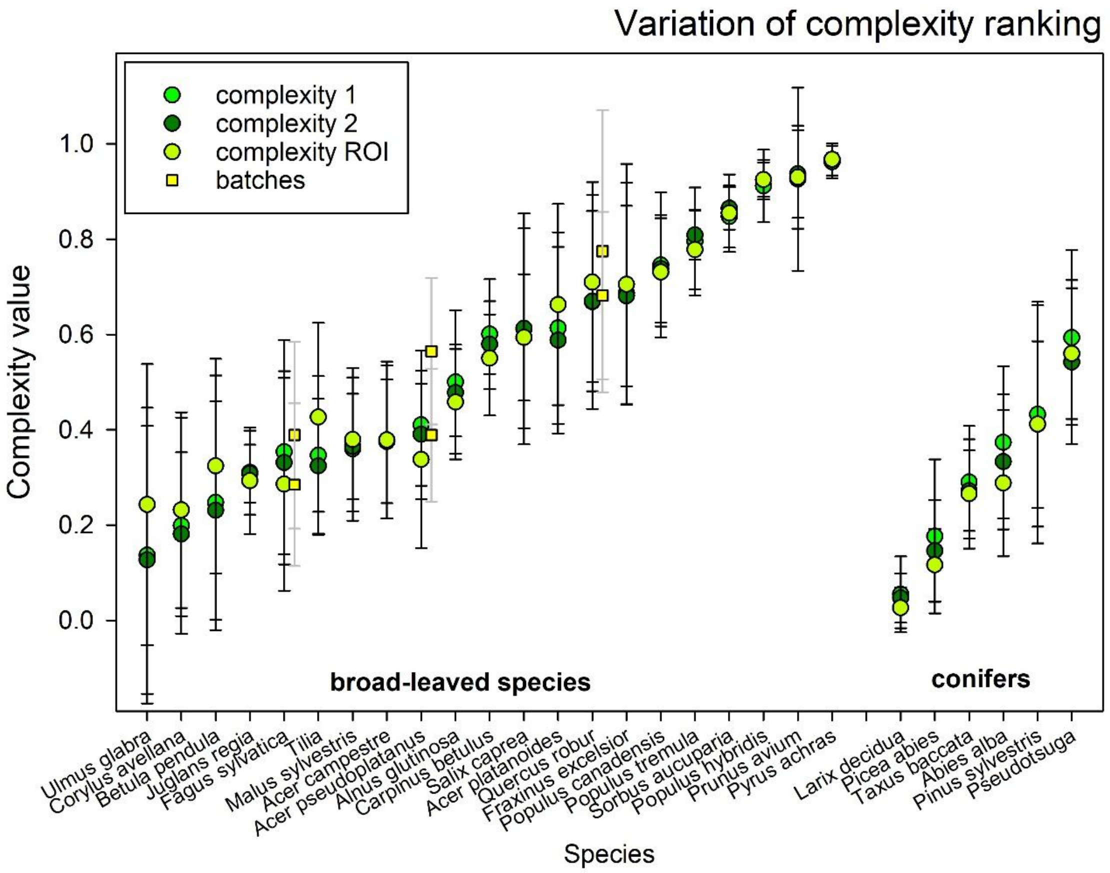

**Supplementary Figure S3**: **Mapping of leaf surface texture complexity groups onto a phylogeny of tree species.** The phylogeny was reconstructed based on published phylogeny of vascular plants (Durka and Michalski, 2012).

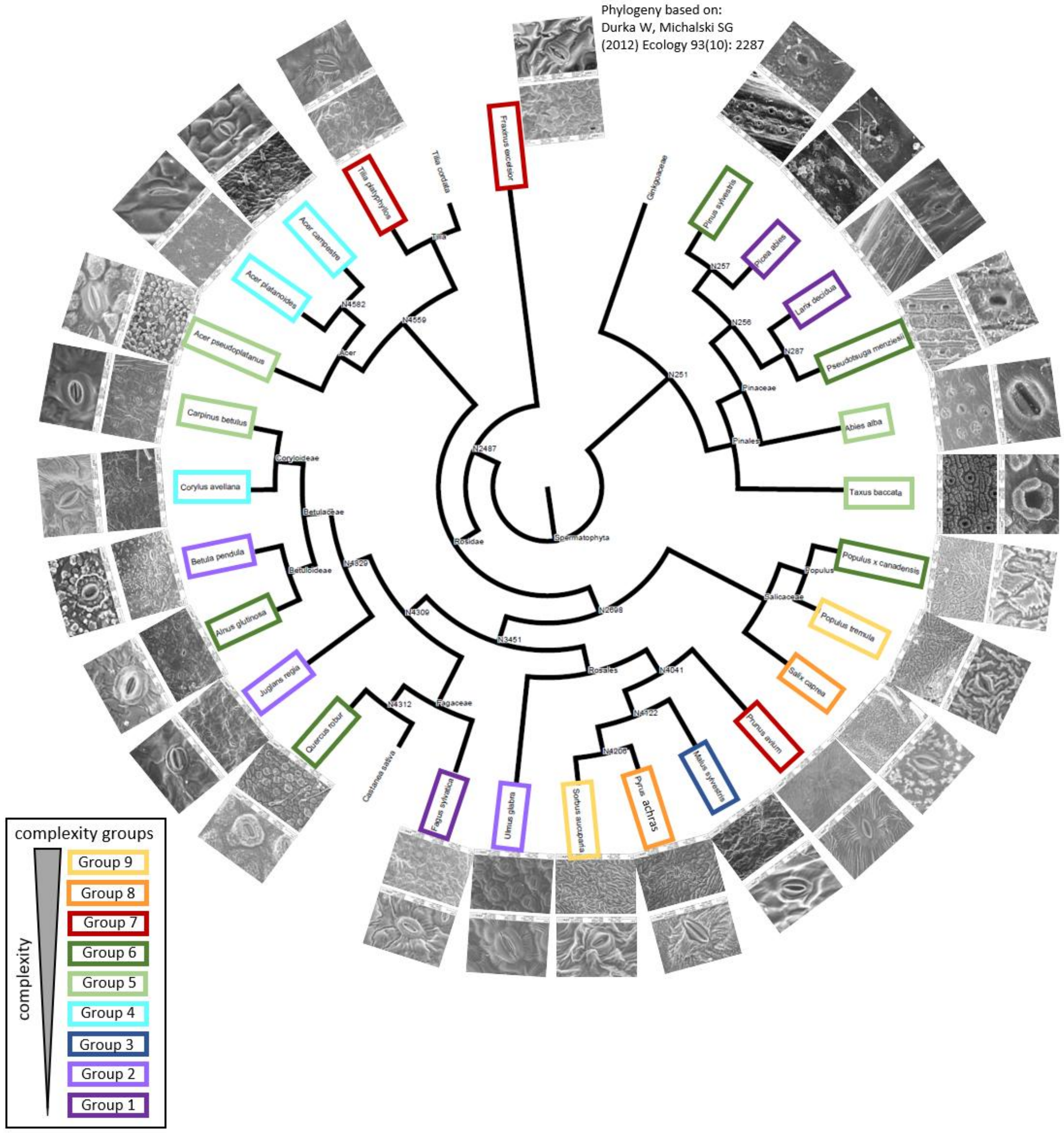

**Supplementary Figure S4**: **Relationship of anatomic and environmental factors with leaf surface complexity.** (**A**) Correlations of leaf surface texture complexity with angle, and stomata morphology. (**B**) Relationship of Ellenberg indicator values of growth habitat characteristics (moisture indicator**,** light indicator, temperature indicator) with leaf surface texture complexity. Indicator values were obtained from published data sets (Tichy *et al.*, 2022). Data points represent averages with standard deviation.

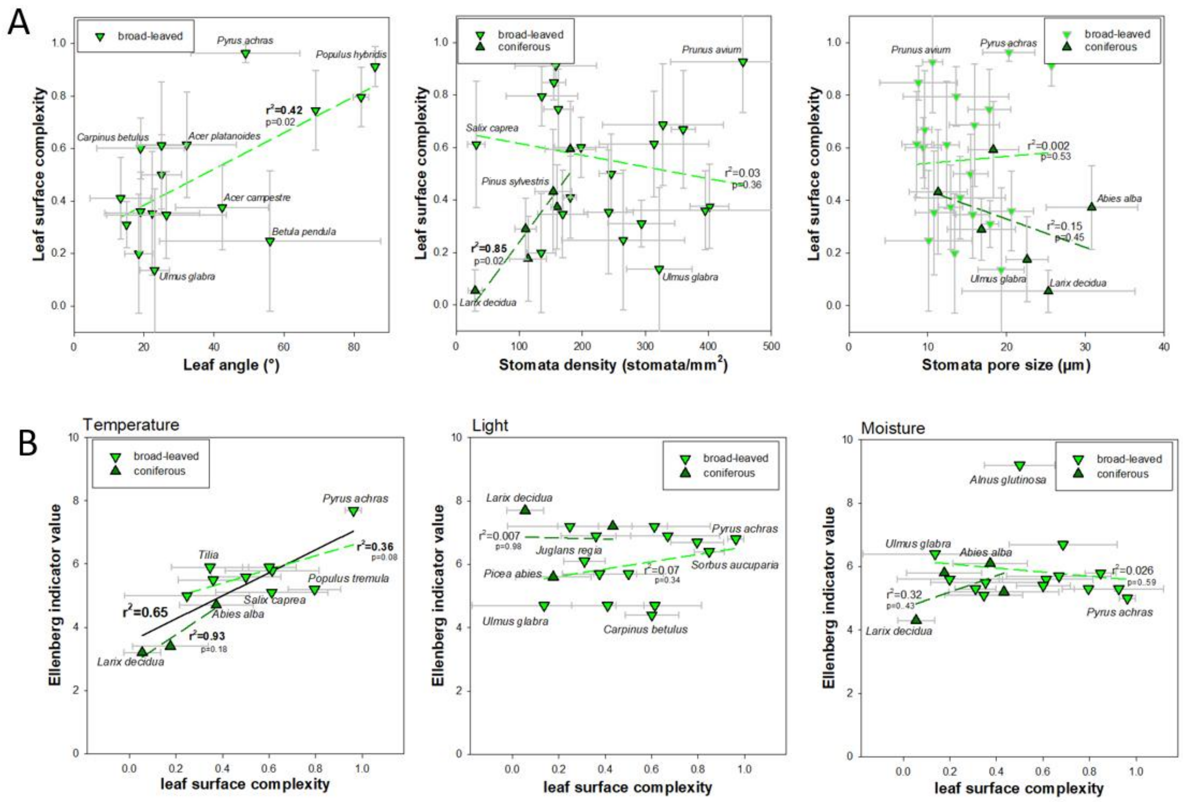

**Supplementary Figure S5**: **Relationship of fungal and bacterial richness with leaf surface texture complexity.** (**A**) Relationship of richness of epiphyllic Basidiomycetae and Ascomycetae with leaf surface texture complexity. (**B**) Relationship of richness of epiphyllic bacteria with leaf surface texture complexity (**C**) Correlation of fungal and bacterial generalists with leaf surface complexity. Data points represent averages with standard deviation. Species richness was assessed by amplified sequence variants (ASV).

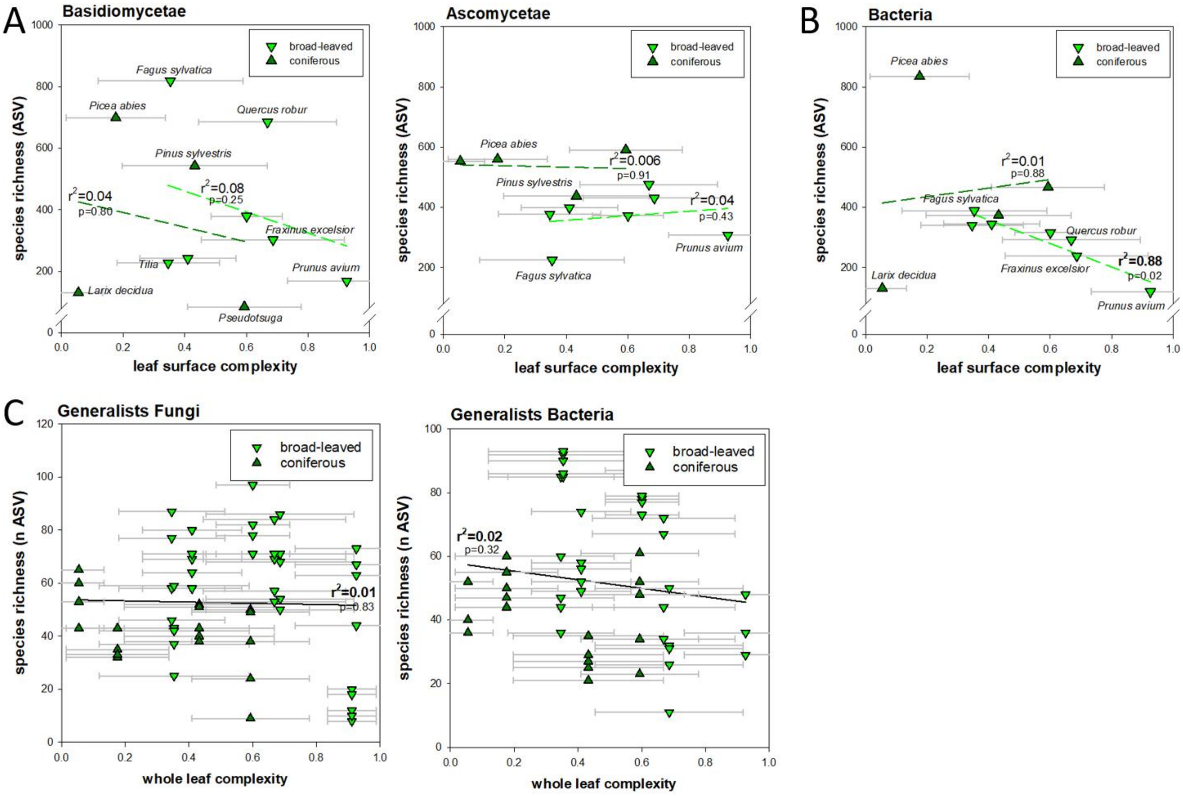

**Supplementary Figure S6**: **Leaf surfaces are poor in nitrogen, as exemplified by surface element analysis.** (**A**) *Quercus robur*. (**B**) *Abies alba,* 2 years. (**C**) *Abies alba*, 4 years. (***D***) *Abies alba*, 8 years. Scanning electron microscopy image (EVO15, Carl Zeiss Microscopy GmbH), and false-color images of elemental abundance (weight%) as determined Energy Dispersive X-ray Spectroscopy (EDS). Bar graphs show weight% of C, O, and N at areas with hyphae (black) or stomata (blue) and a reference region (white).

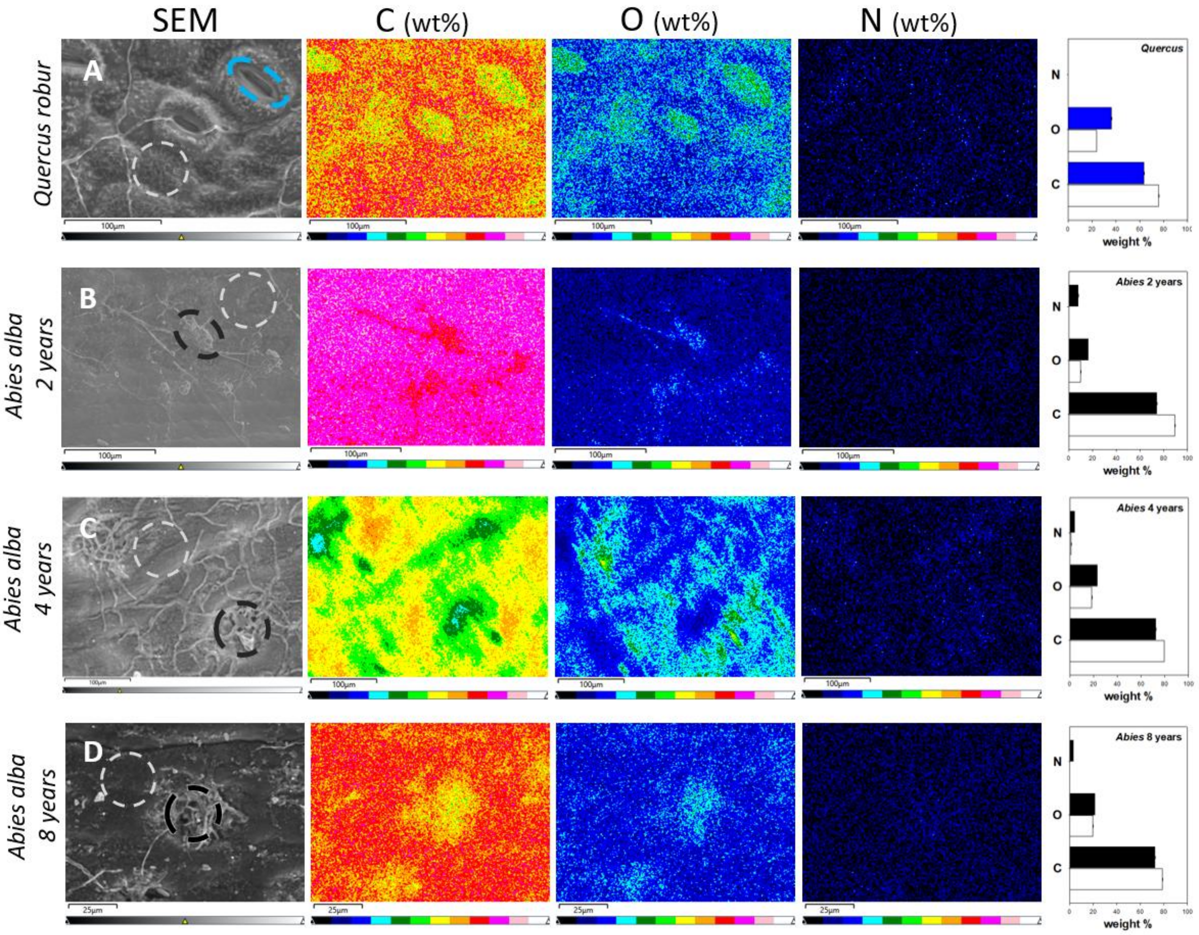
